# Machine learning directed organoid morphogenesis uncovers an excitable system driving human axial elongation

**DOI:** 10.1101/2022.05.10.491358

**Authors:** Giridhar M. Anand, Heitor C. Megale, Sean H. Murphy, Theresa Weis, Zuwan Lin, Yichun He, Xiao Wang, Jia Liu, Sharad Ramanathan

## Abstract

The human embryo breaks symmetry to form the anterior-posterior axis of the body. As the embryo elongates along this axis, progenitors in the tailbud give rise to axial tissues that generate the spinal cord, skeleton, and musculature. The mechanisms underlying human axial elongation are unknown. While ethics necessitate *in vitro* studies, the variability of human organoid systems has hindered mechanistic insights. Here we developed a bioengineering and machine learning framework that optimizes symmetry breaking by tuning the spatial coupling between human pluripotent stem cell-derived organoids. This framework enabled the reproducible generation of hundreds of axially elongating organoids, each possessing a tailbud and an epithelial neural tube with a single lumen. We discovered that an excitable system composed of WNT and FGF signaling drives axial elongation through the induction of a signaling center in the form of neuromesodermal progenitor (NMP)-like cells. The ability of NMP-like cells to function as a signaling center and drive elongation is independent of their potency to generate mesodermal cell types. We further discovered that the instability of the underlying excitable system is suppressed by secreted WNT inhibitors of the secreted frizzled-related protein (SFRP) family. Absence of these inhibitors led to the formation of ectopic tailbuds and branches. Our results identify mechanisms governing stable human axial elongation to achieve robust morphogenesis.

## Introduction

During development, the human embryo elongates along its anterior-posterior (A-P) axis (Ferrer-Vaquer and Hadjantonakis, 2013). Over the course of elongation, axial progenitor cells specified in the tailbud of the embryo give rise to the posterior neural tube and paraxial mesoderm (Anderson et al., 2013; Coutaud and Pilon, 2013; Rodrigo Albors et al., 2018; Wymeersch et al., 2016; Wymeersch et al., 2021). While the signals involved in A-P axis specification have been studied extensively, the roles and relationships of cell types and signals driving elongation are poorly understood, especially in humans (Binagui-Casas et al., 2021). Furthermore, the mechanisms that govern stable elongation along a single axis are unknown.

Elucidating the mechanisms underlying human axial elongation requires the use of *in vitro* stem cell systems. In multiple instances, both human and mouse pluripotent stem cell-derived organoids show spontaneous symmetry breaking when exposed to external signals and generate patterned tissues similar to those in developing mammalian embryos (Moris et al., 2020; van den Brink et al., 2020; Veenvliet et al., 2020). However, the inherent variability in such *in vitro* systems makes it challenging to perform mechanistic studies (Gupta et al., 2021; van den Brink and van Oudenaarden, 2021). Overcoming such stochasticity is therefore essential to achieve the goal of understanding human axial elongation.

Here, we developed an approach that combines bioengineering with machine learning to learn to build hundreds of organoids that reproducibly undergo deterministic A-P symmetry breaking and axial elongation. Using a combination of single cell and *in situ* sequencing, live imaging, and perturbations, we characterized elongating organoids and uncovered distinct roles for FGF and WNT signaling during elongation. By determining conditions that sustain organoid elongation in the absence of external signals, we discovered that elongation is driven by an excitable system mediated by neuromesodermal progenitor (NMP)-like cells. We further identified a role for secreted inhibitors in maintaining the stability of this excitable system. Finally, we discuss the implications of our findings for modeling related morphogenetic processes and more broadly for building robust *in vitro* models of human development.

## Results

### Spatial coupling of organoids influences their axial patterning and morphogenesis

In physical systems that undergo symmetry breaking, the underlying degrees of freedom can be coupled to reduce the entropy of the ground state (Anderson, 1984). Furthermore, in such systems, a desired ground state can be achieved by learning the correct couplings (Hopfield, 1982). Inspired by these results, we asked whether we could couple organoids to increase the robustness of symmetry breaking and achieve the desired patterning and morphogenesis accompanying axial elongation *in vivo*. Specifically, we sought to build organoids that break A-P symmetry, specify a posterior tailbud and anterior neural tube, and elongate along the A-P axis. (Fig. 1A). To test whether organoids could be coupled to each other, we cultured human pluripotent stem cells (hPSCs) on coverslips with randomly positioned micropatterns of extracellular matrix (see Methods). Upon addition of matrix to the culture medium, hPSCs on each micropattern formed an epithelial cyst enclosing a single lumen, closely approximating the epiblast *in vivo* (Zheng et al., 2019) (Fig. 1B). We obtained distinct random arrangements of epithelial cysts on each coverslip, each with a different average density (number of cysts per unit area). To generate organoids, we exposed these cysts to conditions that confer a posterior epiblast identity (Wilson et al., 2009; Wymeersch *et al*., 2021) by activating the canonical WNT pathway using the GSK3ß inhibitor CHIR99021 (CHIR), while inhibiting the BMP and TGFß pathways using the SMAD inhibitors LDN193189 (LDN) and A83-01 respectively (Fig. 1C). After four days of differentiation, we stained the differentiated organoids for CDX2, an axial progenitor marker (Cambray and Wilson, 2007; van den Akker et al., 2002) and SOX1, a neural tube marker (Pevny et al., 1998). Under identical differentiation conditions, the proportion of SOX1+ versus CDX2+ fates on a coverslip was positively correlated with average organoid density (r = 0.98) (Fig. 1D, Fig. S1, A and B). Furthermore, organoids with polarized expression of SOX1 and CDX2 tended to elongate along the SOX1-CDX2 axis, while unpolarized organoids did not elongate (Fig. 1D, right). To quantify the variability in polarization across organoids, we defined polarization μ as the distance between the centroid of CDX2+ cells and the centroid of the organoid (see Methods). On individual coverslips, we generally observed large variations in polarization and axial elongation. However, in replicate experiments with identical initial arrangements, organoids in the same position on the micropatterns showed lower variability in polarization and elongation (Fig. S1, C and D). We concluded that organoids could be coupled and could influence each other’s patterning and morphogenesis based on their relative spatial arrangement.

**Fig. 1.**
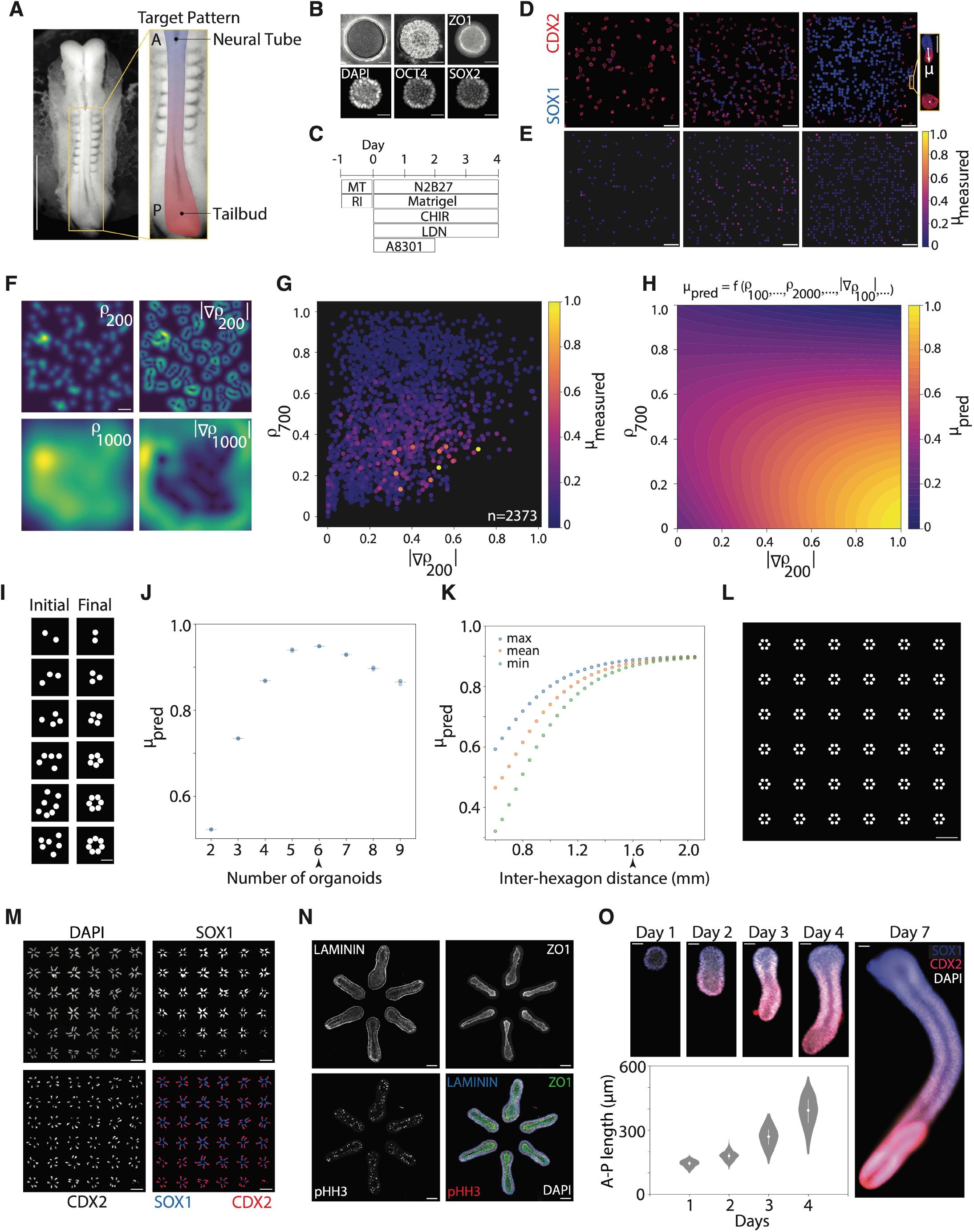
Machine learning-driven optimization of organoid arrangements enables reproducible axial elongation. **(A)** *Left:* Micrograph of a human embryo at Carnegie Stage 10 from the Kyoto Collection. *Right:* Zoomed-in view of posterior neural tube, pseudo-colored to highlight putative anterior neural tube (blue) and posterior tailbud (red). Scale bar 1 mm. **(B)** *Top left:* Phase contrast image of 150 μm diameter circular PDMS stamp. *Top middle:* Phase contrast image of micropatterned pluripotent epithelial cyst on day 1. *Top right:* Cyst stained for tight junction marker ZO-1. *Bottom*: Cyst stained for nuclear markers DAPI, OCT4, and SOX2. Scale bar 50 μm. **(C)** Organoid differentiation protocol: day -1, hPSCs seeded in mTeSR Plus (MT) with ROCK inhibitor Y-27632 (RI); day 0, media changed to N2B27 with Matrigel, CHIR99021 (CHIR), LDN193189 (LDN), and A83-01; day 2, media changed to N2B27 with CHIR and LDN. Matrigel formed a gel that was maintained throughout 4 days of differentiation. **(D)** *Left:* Organoids from low density (top), medium density (middle), and high density (bottom) random arrays on day 4, stained for SOX1 and CDX2. Scale bar 1 mm. *Bottom left:* Zoomed-in view of an unpolarized CDX2+ organoid (bottom) and polarized organoid elongated along the SOX1-CDX2 axis (top). μ measures the anterior-posterior (A-P) polarization of CDX2 expression (see Methods). Scale bar 200 μm. **(E)** Measured polarization (μ_measured_) of organoids in (D) plotted on color scale at corresponding micropattern locations in the array. **(F)** *Left:* Heat map of organoid density on random arrays evaluated with a Gaussian filter at length scales 3 = 200 μm (top) and 3 = 1000 μm (bottom). *Right*: Visualization of gradient of organoid density using a Sobel filter, evaluated at length scales 3 = 200 μm (top) and 3 = 1000 μm (bottom). For gradient calculations, see Supplementary Methods. Scale bar 1 mm. (**G)** μ_measured_ of each organoid (n=2373) from 12 random arrays as a function of selected features, gradient of organoid density at 3 = 200 μm (|∇ρ_200_|) and density at 3 = 700 (ρ_700_). **(H)** Predicted polarization (μ_pred_) for organoids learned using kernel ridge regression with a radial basis function kernel from data in (G) (see Supplementary Methods), plotted as a function of |∇ρ_200_| and ρ_700_. **(I)** Initial (left) and final (right) timepoint images of simulated annealing on organoid locations using the function learned in (H) with the number of organoids ranging from 2 (top) to 7 (bottom). Scale bar 300 μm. **(J)**, μ_pred_ as a function of number of organoids, with a maximum for 6 organoids arranged in a hexagon. (**K)** maximum, mean, and minimum values of μ_pred_ for hexagonally arranged organoids as a function of inter-hexagon distance. Arrowhead indicates distance selected for experiments (1.6 mm). **(L)** Image of computationally optimized hexagonal array. Scale bar 1 mm. **(M)** Organoids on the optimized hexagonal array on day 4, stained for DAPI, SOX1, and CDX2. All organoids that correctly formed a cyst within a hexagonal unit on day 1 possessed polarized CDX2+ and SOX1+ domains and elongated (n=204). Scale bar 1 mm. **(N)**, *Top: R*epresentative organoids from hexagonal array on consecutive days stained for DAPI, SOX1, and CDX2. Scale bar 50 μm. *Bottom:* Measured A-P lengths of organoids from hexagonal array over time (n=216), based on maximum Feret diameter. **(O)** Confocal images of organoids from hexagonal array on day 4 stained for laminin subunit gamma-1 (laminin), tight junction protein 1 (ZO1), and phospho-histone H3 (pHH3). Organoids display apicobasal polarity with apical mitosis and a basement membrane. Scale bar 100 μm.

### Machine learning driven optimization of organoid arrangements enables reproducible axial elongation

To systematically learn how to arrange organoids to obtain the desired patterning and morphogenesis accompanying axial elongation, we used approaches from machine learning (Friedman et al., 2001). We first measured the polarization of CDX2 expression 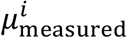 for each organoid *i* in all 2,373 organoids across our random micropattern experiments (Fig. 1E). We then calculated a multi-dimensional feature vector for each organoid, 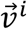, with components consisting of the organoid density *ρ*^*i*^ and the gradient of organoid density |∇*ρ*^*i*^| over a range of length scales from 100 μm to 2000 μm, *i*.*e*., 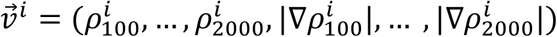 (Fig. 1F). We sought to learn the function *f* of the feature vector that predicted the polarization of each organoid *i, i*.*e*., 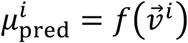, with the minimum least squares error, 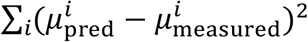. We used sequential forward selection to determine the most predictive subset of features in 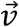, followed by kernel ridge regression with a radial basis function kernel (Luo et al., 2013) and five-fold cross validation to learn the parameters of *f*. We determined that two components of 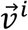 were most predictive of organoid polarization: the organoid density at 700 μm, 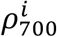, and the gradient of organoid density at 200 μm, 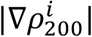 (Fig. 1G). Using the learned function *f*, we mapped out the predicted polarization of organoids in the space of these two components (Fig. 1H). We then performed simulations to modify the arrangements of organoids until each organoid was predicted to have maximum polarization (see Supplementary Methods). To do so, we first performed simulated annealing on small numbers of organoids ranging from 2 to 9, and found that 6 organoids arranged in a hexagon achieved the maximum predicted polarization (Fig. 1, I and J). To determine the arrangement of these hexagonal units on a coverslip, we calculated *μ*_pred_ for each organoid as a function of the distance between these units. We selected the distance such that the predicted polarization was uniformly high across all organoids while fitting as many organoids as possible on a single coverslip (Fig. 1K). The computationally selected arrangement thus consisted of a 6 by 6 array of hexagonal units spaced 1.6 mm apart (Fig. 1L). We conducted our differentiation experiments on coverslips micropatterned with this hexagonal array. After 4 days, every organoid in the array that initially formed a cyst within a hexagonal unit developed a CDX2+ tailbud domain, a SOX1+ neural tube domain, and elongated axially (n=204) (Fig. 1M). Thus, we were able to tune the coupling between organoids by optimizing their spatial arrangement, and induce them to jointly break A-P symmetry to develop into our desired target pattern.

We further found that the organoids recapitulated several aspects of the elongating neural tube *in vivo*. In particular, elongating organoids contained a single lumen enclosed by apical tight junctions (marked by ZO1), a basement membrane (marked by laminin), and exhibited apical mitosis (marked by phospho-histone H3) consistent with interkinetic nuclear migration (Fig. 1N). We also observed pseudostratified SOX2+ nuclei as well as junctional N-cadherin and apical F-actin (Fig. S1E). Upon staining organoids for CDX2 and SOX1 at sequential timepoints, we found that CDX2+ axial progenitors were specified at the prospective posterior pole by day 2 of differentiation and maintained throughout elongation, and a SOX1+ neural tube was specified anteriorly that increased in length over time (Fig. 1O, top and Fig. S1, F and G). The median A-P axis length of organoids increased from approximately 150 μm on day 1 to approximately 400 μm on day 4 (Fig. 1O, bottom and Fig. S1, H and I), and continued to increase through day 7, by which time organoids reached lengths over 1 mm (Fig. 1O, right).

### Elongating organoids contain hindbrain, spinal cord, and axial progenitors

To identify the cell types present in elongating organoids, we used single-cell RNA sequencing (scRNA-seq). We performed scRNA-seq on cells pooled from 12 organoids on day 4 of differentiation to obtain high-quality transcriptomes of 7,752 cells. Using unsupervised sparse multimodal decomposition (SMD) (Melton and Ramanathan, 2021) to simultaneously infer the identity of key genes and cell types, we identified a subspace of 26 genes in which we could cluster cells into 4 anterior and posterior progenitor cell types, corresponding to spatially distinct regions along the A-P axis. The observed cell types included axial progenitors (marked by CDX2) as well as progenitors of the cervical spinal cord (marked by HOXB4 and CRABP1), posterior hindbrain (marked by NR2F1), and anterior hindbrain (marked by EGR2). We also observed a small cluster of neuronal cells (marked by HES6 and BTG2) (Fig. 2, A and B, Fig. S2A). Interestingly, we did not observe any transcriptional signatures of neuromesodermal progenitors (NMPs, marked by TBXT and SOX2), paraxial mesoderm derivatives (marked by TBX6 and MEOX1), or lateral mesoderm derivatives (marked by HAND2 and OSR1) (Fig. S2C), suggesting that elongation can occur even in the absence of these cell types.

**Fig. 2.**
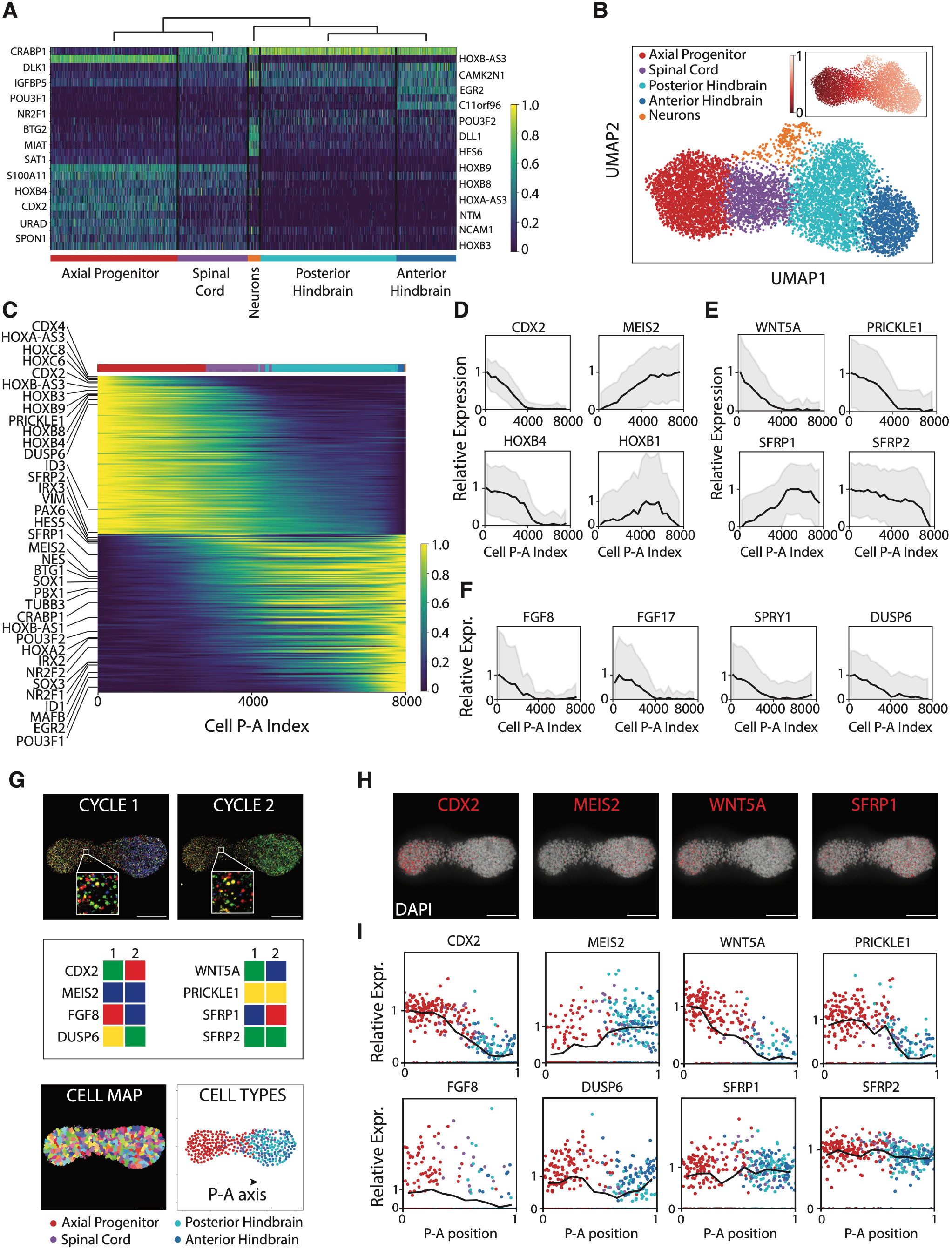
Single cell sequencing identifies posteriorly polarized WNT and FGF/ERK signaling in axially elongating organoids. **(A)** Gene expression heatmap of cells (n=7752) from elongating organoids on day 4 using single cell RNA-sequencing (scRNA-seq). Cells clustered with the Louvain method in a 26-dimensional gene space selected by sparse multimodal decomposition (see Methods) show axial progenitor (CDX2+), spinal cord (HOXB4+ CRABP1+), posterior hindbrain (NR2F1+), anterior hindbrain (EGR2+), and neuronal (HES6+ BTG2+) identities. Expression values are log- and min/max-normalized across cells. **(B)** UMAP plot of cells from scRNA-seq colored by cell type (bottom) and posterior anterior (P-A) pseudo-spatial coordinate obtained by diffusion mapping (inset). **(C)** Gene expression heatmap of top 300 differentially expressed genes ordered by computationally inferred cell P-A index. Expression values are log- and min/max-normalized across cells. **(D-F)** Gene expression profiles as a function of cell P-A index for **(E)** transcription factors CDX2, MEIS2, HOXB4, HOXB1, **(E)** non-canonical WNT ligand WNT5A and non-canonical WNT pathway component PRICKLE1, secreted WNT inhibitors SFRP1, SFRP2, (**F**) FGF ligands FGF8 and FGF17, FGF/ERK pathway targets SPRY1, DUSP6. Each plot displays mean (black) and standard deviation (gray) of log-normalized relative expression values partitioned into 20 bins of equal size. **(G)** *Top: In situ* sequencing (STARmap) of selected genes in organoid sample on day 4. Each gene transcript is encoded by a unique 2-color sequence across 2 cycles of sequencing (see legend). *Bottom*: Using locations of nuclei, transcripts were mapped to individual cells (left). This map was used to assign cell types (right) corresponding to **(A)**. Scale bar: 100 μm. **(H)** Decoded transcripts in red for selected genes CDX2, MEIS2, WNT5A, SFRP1, overlaid on nuclei (DAPI). Scale bar: 100 μm. **(I)** Spatial P-A gene expression profiles for selected genes in **(D-F)** colored by cell type. Each plot displays mean (black) and per-cell (points) log-normalized relative expression values partitioned 10 bins of equal length.

### WNT and FGF signals are posteriorly polarized in elongating organoids

Having identified the constituent cell types, we proceeded to investigate the underlying signals that initiate and drive axial elongation. To infer spatial profiles of signaling pathways in organoids, we built a diffusion map (Haghverdi et al., 2016) in the Euclidean space of genes identified by SMD to order cells from scRNA-seq along a putative posterior-anterior (P-A) axis (Fig. 2, B and C, Fig. S2, C and D). We first validated this pseudo-spatial ordering by verifying the expected P-A localization of axial progenitor, spinal cord, and hindbrain transcription factors such as CDX2, MEIS2, HOXB4, and HOXB1 (Fig. 2D). We then used this pseudo-spatial P-A axis to determine profiles of signaling pathways implicated in axis formation. While we did not detect the expression of any canonical WNT ligands, we observed posteriorly polarized expression of noncanonical WNT ligands WNT5A and WNT5B as well as noncanonical WNT component PRICKLE1, and global expression of secreted WNT inhibitors SFRP1 and SFRP2 (Fig. 2E, Fig. S2B). We also observed posteriorly polarized expression of FGF ligands FGF8 and FGF17, and FGF/ERK targets SPRY1 and DUSP6 (Fig. 2F, Fig. S2B). Consistent with their inferred polarization, WNT and FGF ligands were also identified by an unbiased search for differentially expressed secreted factors across progenitor clusters (Fig. S2C). To validate these signaling profiles, we performed *in situ* sequencing on day 4 organoids using STARmap (Wang et al., 2018) (Fig. 2G). To do so, we implemented a revised STARmap protocol and corresponding computational pipeline compatible with 3D tissue sequencing and analysis to spatially map 16 genes selected from scRNA-seq data (see Supplementary Methods). Across three replicates, we observed posteriorly polarized expression of CDX2, WNT5A, PRICKLE1, and FGF8, and global expression of SFRP1 and SFRP2, matching our inferences from scRNA-seq data (Fig. 2, H and I, Fig. S2F). The results from *in situ* sequencing validated the cell types and diffusion map trajectories inferred from scRNA-seq (Fig S2, G to I). Our analyses thus demonstrate that WNT and FGF/ERK signals are localized posteriorly to the tailbud of axially elongating organoids.

### WNT ligands drive axial elongation downstream of canonical WNT activity

Given that WNT signals are polarized in organoids, we next asked whether WNT ligands drive axial elongation downstream of CHIR-mediated canonical WNT activation. We inhibited WNT ligand activity using the Porcupine inhibitor IWP3, which blocks WNT secretion by preventing its palmitoylation (Chen et al., 2009). We found that IWP3-treated organoids failed to elongate, and retained a spherical shape after four days of differentiation (Fig. 3, A and B, Fig. S3A). However, IWP3-treated organoids retained a significant posterior CDX2+ population, similar in size to control organoids when adjusted for A-P length (Fig. 3C, Fig. S3, A and B). This suggested that in the presence of CHIR, secreted WNT ligands are necessary for elongation, but not for the initial specification of posterior identity. To test whether secreted WNT ligands were required continuously for elongation or act as an initial trigger, we treated elongating organoids on day 2 with IWP3. We found that organoids were truncated by day 4 despite maintenance of CDX2+ domains (Fig. 3, D to F, Fig. S3B), demonstrating that continuous secretion of WNT ligands was necessary for elongation. To explore the role of WNT ligands in regulating cell behavior during elongation, we performed live confocal time-lapse imaging of organoids expressing ZO1-EGFP, a fusion reporter of apical tight junctions, with or without IWP3 treatment on day 2 of differentiation. We verified that IWP3 treatment led to truncation of ZO1-EGFP+ organoids even in the presence of a posterior CDX2+ domain (Fig. 3G, Fig. S3G). By tracking the displacement of individual cell clones over the course of fourteen hours (Fig. 3H, Movie S1), we found that neighboring clones in control organoids exhibited higher mean squared relative displacements compared to IWP3-treated organoids (Fig. 3I). This suggested that cells experienced increased fluidity (Mongera et al., 2018) in the presence of WNT ligands. We also found that control organoids displayed directed cell movements in the plane of the epithelium along the direction of the elongation axis. In contrast, cell movements in IWP3-treated organoids were undirected with respect to the elongation axis (Fig. 3J, Fig. S3H). We additionally performed time lapse imaging of organoids expressing H2B-mCherry under control conditions to determine whether cell divisions were oriented with respect to the elongation axis. However, there was no correlation between the angle of cell division and elongation direction (Fig. S3I). We concluded that WNT ligands are continuously required to maintain the fluidity of the epithelium to facilitate rearrangements of neighboring cells in the direction of the elongating axis. Furthermore, the detection of only noncanonical WNT ligands in single cell expression data (Fig. 3E) and the observation that addition of DKK1, a canonical WNT receptor inhibitor, has no effect on elongation (Fig. S3, E and F), support a potential role for noncanonical WNT in driving elongation.

**Fig. 3.**
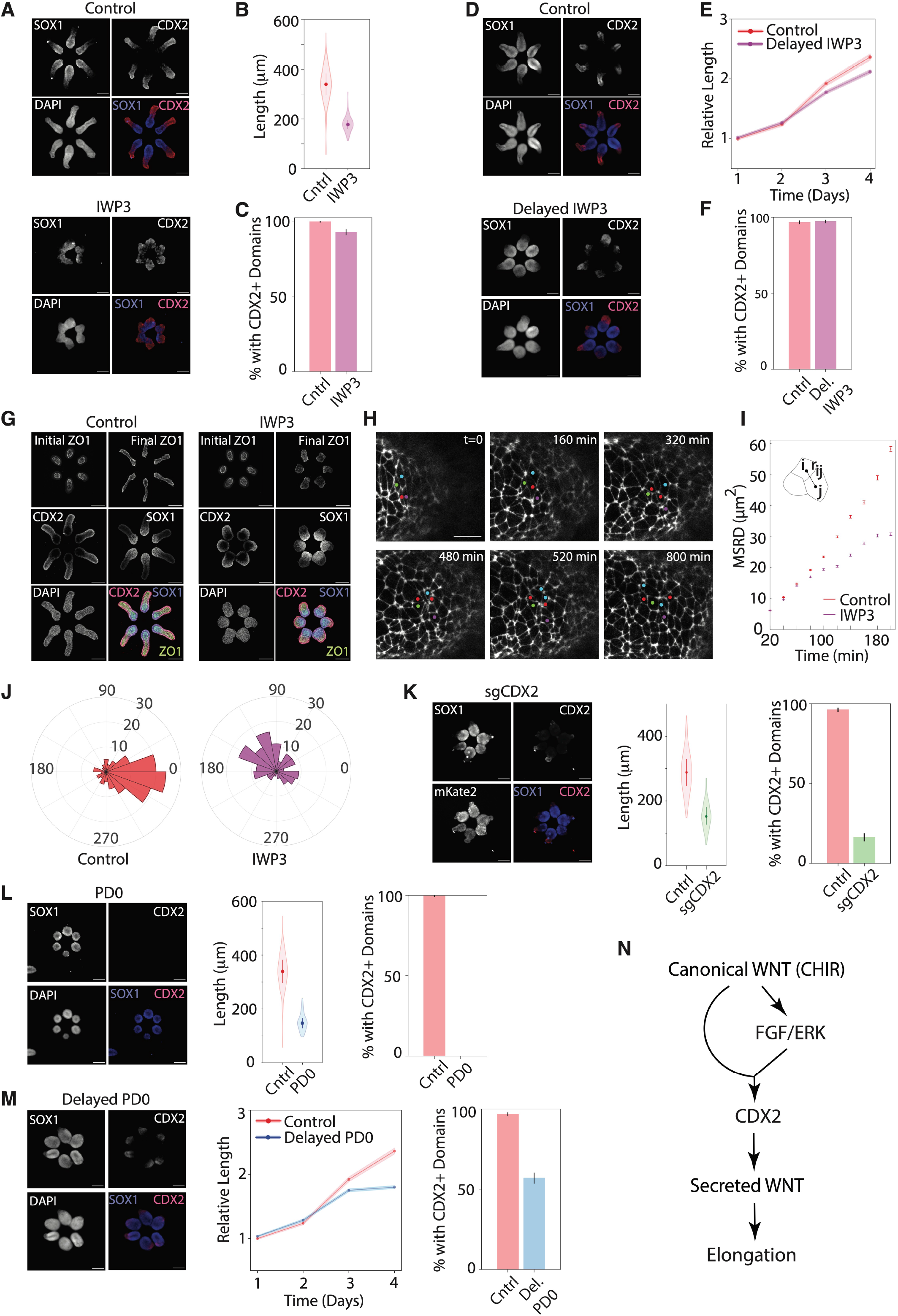
WNT ligands drive elongation downstream of FGF/ERK-dependent CDX2 expression. **(A)** Control (left) and IWP3-treated (right) organoids on day 4 stained for SOX1, CDX2, and DAPI. Scale bar: 200 μm. **(B)** Violin plot of day 4 lengths of control (red) and IWP3-treated organoids (purple) with median (points) and interquartile range (lines) (Control, n=211, IWP3, n=176). **(C)** Bar plot showing percentage of control and IWP3-treated organoids with a CDX2+ domain (see Methods). Error bars denote SEM (Control, n=211, IWP3, n=176). **(D)** Control (left) and 48h-delayed IWP3-treated (right) organoids on day 4 stained for SOX1, CDX2, and DAPI. Scale bar: 200 μm. **(E)** Line plot of relative lengths of control (red) and 48h-delayed IWP3-treated (purple) organoids over time. Shaded regions denote SEM (Control, n=212, IWP3, n=214). **(F)** Bar plot showing percentage of control and 48h-delayed IWP3-treated organoids with a CDX2+ domain. Error bars denote SEM (Control, n=212, IWP3, n=214). **(G)** *First row:* Initial (day 2) and final (day 4) control (left) and IWP3-treated (right) ZO1-EGFP+ organoids. *Second, third row:* ZO1-EGFP+ organoids on day 4 without (left) and with (right) IWP3 treatment, stained for CDX2, SOX1, and DAPI. Scale bar: 200 μm. **(H)** Representative live fluorescence images of tracked ZO1-EGFP+ cells from a control organoid during a 14h time-lapse. Each clone from t=0 is marked with a single color. Scale bar 50 μm. **(I)** Mean squared relative displacement (MSRD) over time of neighboring cells in control (red) and IWP3-treated (purple) organoids. **(J)** Angle histogram of net cell displacements relative to the axis of elongation in control (left, red) and IWP3-treated (right, purple) organoids (Control, n=131, IWP3, n=98). **(K)** *Left:* CRISPRi organoids containing an mKate2-expressing guide RNA construct against CDX2 (sgCDX2) on day 4, stained for SOX1 and CDX2. Scale bar: 200 μm. *Middle*: Violin plot of day 4 lengths of control sgRNA (red) and sgCDX2 (green) CRISPRi organoids (Control, n=213, sgCDX2, n=214). *Right:* Bar plot showing percentage of control (red) and sgCDX2 (green) CRISPRi organoids with a CDX2+ domain. Error bars denote SEM (Control, n=213, sgCDX2, n=214). **(L)** *Left:* PD0-treated organoids on day 4, stained for SOX1, CDX2, and DAPI. Scale bar: 200 μm. *Middle*: Violin plot of day 4 lengths of control (red) and PD0-treated (blue) organoids (Control, n=211, PD0, n=157). *Right:* Bar plot showing percentage of control (red) and PD0-treated (blue) organoids with a CDX2+ domain. Error bars denote SEM (Control, n=211, PD0, n=157). **(M)** *Left:* 48h-delayed PD0-treated organoids on day 4, stained for SOX1, CDX2, and DAPI. Scale bar: 200 μm. *Middle*: Line plot of relative lengths of control (red) and 48h-delayed PD0-treated (purple) organoids over time. Shaded regions denote SEM (Control, n=212, PD0, n=211). *Right:* Bar plot showing percentage of control (left, red) and 48h-delayed PD0-treated (right, blue) organoids with a CDX2+ domain. Error bars denote SEM (Control, n=212, PD0, n=211). **(N)** Signaling model of axial elongation of the neural tube. CHIR-mediated canonical WNT induces FGF/ERK activity, together inducing CDX2 expression. CDX2 promotes expression of secreted WNT ligands, which drive elongation.

### FGF/ERK-dependent CDX2 expression is required for elongation

We next tested the roles of intrinsic and extrinsic signaling factors implicated upstream of WNT ligand activity in the tailbud. Across vertebrates, WNT ligands are induced by CDX transcription factors (Chawengsaksophak et al., 2004). We used CRISPR interference (CRISPRi) to knock down CDX2 expression in a constitutive CRISPRi hPSC line (Mandegar et al., 2016) (see Methods). We generated organoids from CRISPRi lines with an integrated CDX2 guide RNA (gRNA) or nonspecific control gRNA. By day 4, organoids with a control gRNA elongated normally, whereas organoids with a CDX2 gRNA were truncated and lacked CDX2+ domains (Fig. 3K, Fig. S3, J to L), suggesting that CDX2 is essential for organoid elongation. CDX transcription factors are thought to lie downstream of FGF/ERK and canonical WNT signaling in vertebrates (Pownall et al., 1996). To test the role of FGF/ERK, we treated organoids with PD0325901 (PD0), a small molecule inhibitor of ERK1/2 phosphorylation. Following continuous FGF/ERK inhibition, organoids failed to elongate or produce any CDX2+ axial progenitors (Fig. 3L, Fig. S3A). This demonstrated that FGF/ERK was necessary for the induction of CDX2 expression and subsequent axis elongation. To determine whether FGF/ERK signaling was required throughout the elongation process or only at the outset, we inhibited FGF/ERK on day 2 of differentiation. By day 4, organoids were truncated relative to control organoids and downregulated CDX2 expression, showing FGF/ERK was required not only to induce CDX2 but also to maintain its expression level for elongation to occur (Fig. 3M, Fig. S3B). Together, our results support a model in which canonical WNT activity (driven by CHIR) together with FGF/ERK activity (driven by endogenous FGF ligands) induces and maintains CDX2 expression in the tailbud. In turn, CDX2 drives the secretion of WNT ligands, which are essential for directing elongation by modulating cellular rearrangements (Fig. 3N).

### Elongation is sustained by induction of a neuromesodermal progenitor-like signaling center

In experiments described thus far, we required sustained WNT activation through CHIR to drive elongation. We next asked if organoids had the ability to elongate autonomously even in the absence of this external drive. Axial elongation is thought to be sustained *in vivo* by NMPs, which produce WNT and FGF ligands (Amin et al., 2016). Studies in vertebrates have further demonstrated that canonical WNT and FGF/ERK pathways are engaged in a positive feedback loop in a variety of developmental contexts, including the zebrafish tailbud (Canning et al., 2007; Martin and Kimelman, 2008; Takeuchi et al., 2003). We sought to exploit this feedback between WNT and FGF activity to sustain elongation. Thus, we exposed organoids to a transient 24-hour pulse of recombinant FGF2 prior to withdrawing CHIR on day 2 of differentiation. Whereas under control conditions, CHIR withdrawal led to truncated organoids with reduced CDX2 expression (Fig. 4A, middle, Fig. 4B, Fig. S4, A and B), FGF pulse-treated organoids continued to elongate through day 4 of differentiation, showing that a pulse of FGF was sufficient to sustain elongation (Fig. 4A, bottom, Fig. 4B, Fig. S4, A and B). To verify induction of NMPs, we stained FGF pulse-treated organoids for TBXT and SOX2, transcription factors which are known to jointly mark NMPs *in vivo*. Cells in the tailbud of FGF-treated organoids stained positively at the end of the pulse for TBXT and SOX2 (Fig. 4, C to E, Fig. S4, C and D). In contrast, control organoids exposed to CHIR alone failed to express TBXT. Furthermore, FGF pulse-treated organoids maintained TBXT+ SOX2+ cells in the CDX2+ tailbud region throughout elongation, although these cells decreased in number over time (Fig. S4E). We additionally found that a pulse of CHIR, at a concentration higher than in control conditions, could generate a similar result (Fig. S4, A and B) in an FGF-dependent manner. In vertebrates, WNT and FGF ligand production in NMPs is thought to be driven in part by transcription factor TBXT (Amin et al., 2016). To test the requirement of TBXT for sustained elongation, we generated organoids from CRISPRi lines with an integrated TBXT guide RNA (Fig. S4F), and exposed them to continuous CHIR or FGF pulse conditions. Compared to CRISPRi organoids containing a control gRNA, organoids with a TBXT gRNA were truncated following the FGF pulse (Fig. S4G). This is consistent with the possibility that TBXT is required for production of canonical WNT and FGF/ERK signals by NMPs during axial elongation. Accordingly, TBXT+ SOX2+ cells from axial organoids express WNT3A and WNT8A as well as FGF4 and FGF8, which activate canonical WNT and FGF/ERK pathways, respectively (Yaman et al., 2022). Thus, induction of an NMP-like signaling center enables self-sustained axial elongation of the neural tube, likely through continuous production of posterior signals.

**Fig. 4.**
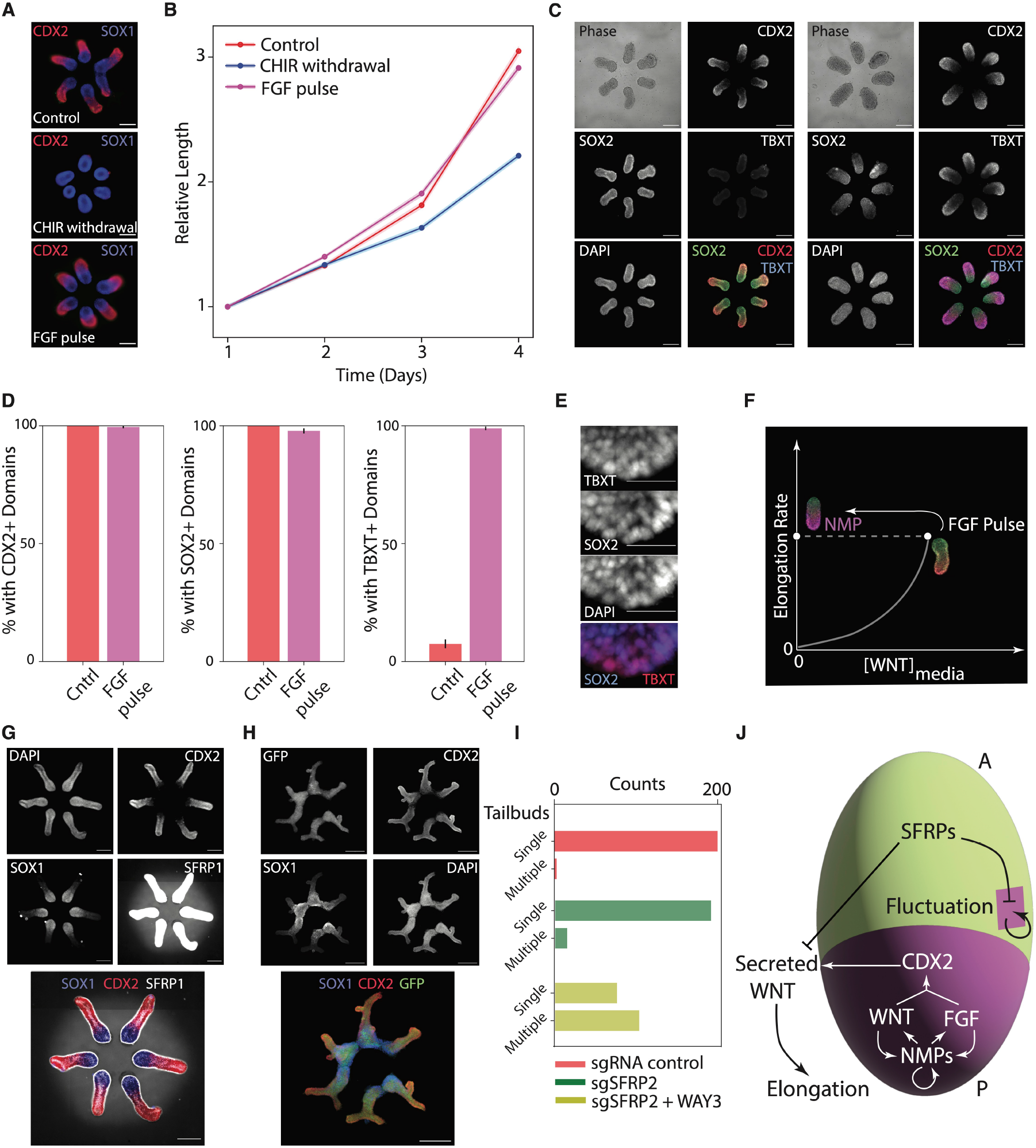
An excitable system driving axial elongation is triggered by induction of an NMP-like signaling center and is stabilized by secreted inhibitors. **(A)** Day 4 control organoids (top), organoids subjected to CHIR withdrawal on day 2 (middle), and organoids treated with an FGF pulse from day 1 to day 2 prior to CHIR withdrawal on day 2 (bottom), stained for CDX2 in red and SOX1 in blue. Scale bar: 200 μm. **(B)** Line plot of relative lengths over time of organoids exposed to control (red), CHIR withdrawal (blue), and FGF pulse (red) conditions. Shaded regions denote SEM (Control, n = 216, CHIR withdrawal, n = 214, FGF pulse, n = 216) **(C)** Phase contrast and fluorescence images of day 2 organoids in control (left) and FGF pulse + CHIR withdrawal (right) conditions, stained for CDX2, SOX2, TBXT, and DAPI. **(D)** Bar plots showing percentage of day 2 organoids in control (red) and FGF pulse + CHIR withdrawal (magenta) conditions with a CDX2+ domain (left), SOX2+ domain (middle), and TBXT+ domain (right). Error bars denote SEM (Control, n = 199, FGF pulse, n = 184). **(E)** Zoomed-in fluorescence images of the posterior side of day 2 organoids after a 24-hour FGF pulse, stained for TBXT, SOX2, and DAPI. Co-expression of nuclear TBXT and SOX2 suggest neuromesodermal progenitor (NMP)-like identity. Scale bar: 100 μm. **(F)** Model of axial elongation as an excitable system. Organoids elongate in media with WNT signal. The rate of elongation is zero when the concentration of WNT in the media, [WNT]_media_, is zero. An FGF pulse induces an NMP-like signaling center at the posterior end of organoids. In this condition, the rate of elongation is sustained when [WNT]_media_ is zero. **(G)** Control organoids on day 4, stained for DAPI, CDX2, SOX1, and SFRP1. Images were contrast-adjusted to visualize extracellular SFRP1. In the merged image, DAPI was subtracted to emphasize extracellular SFRP1. Scale bar: 200 μm. **(H)** Day 4 CRISPRi organoids containing a GFP-expressing guide RNA construct against SFRP2 (sgSFRP2) and treated with WAY-3, stained for CDX2, SOX1, and DAPI. Scale bar 200 μm. **(I)** Bar plots showing number of organoids with single or multiple tails in control (top, red, n = 203), sgSFRP2 (middle, green, n = 206), and WAY3-treated sgSFRP2 (bottom, yellow, n = 179) CRISPRi organoids. **(J)** Proposed mechanisms underlying axial elongation. Low levels of posteriorly localized FGF/ERK and canonical WNT activity induce CDX2 expression, which in turn induces secretion of WNT ligands, driving axial elongation. High levels of FGF/ERK and canonical WNT activity induce NMPs, which continuously produce FGF/ERK and canonical WNT ligands to maintain CDX2 expression. Local fluctuations in WNT ligand levels (purple) are buffered by global secretion of SFRP1/2, ensuring that elongation occurs only along a single A-P axis.

### Secreted WNT inhibitors are required for stable axial elongation

These results suggest that organoids can be excited from a state in which elongation requires external WNT signals to one in which it can be self-sustained (Fig. 4F). Such excitable systems are known to be susceptible to noise-driven instabilities that can trigger runaway events (Murray, 1989). We therefore asked whether inhibitors of WNT or FGF were necessary for stable elongation. In our scRNA-seq and STARmap data, we observed globally high expression levels of secreted WNT inhibitors SFRP1 and SFRP2 (Fig. 2, E and I, Fig. S2, B and C). To verify that SFRPs were being secreted by organoids, we stained organoids for SFRP1 protein on day 4. We found that SFRP1 protein was present extracellularly and was distributed in a gradient along the A-P axis of each organoid (Fig. 4G). To test whether SFRP1 and SFRP2 suppressed runaway events leading to ectopic sites of elongation, we generated organoids from CRISPRi lines with an integrated SFRP2 gRNA, and treated organoids with WAY-316606 (WAY-3), a small molecule which inhibits SFRP1 activity by preventing its interaction with WNT ligands (Bodine et al., 2009). While knockdown of SFRP2 or inhibition of SFRP1 alone did not produce severe defects (Fig. S4H), simultaneous SFRP1 inhibition and SFRP2 knockdown resulted in ectopic tailbud formation and elongation, characterized by branched structures (Fig. 4H) and an increased average number of tailbuds per organoid (Fig. 4I). Thus, redundantly acting WNT inhibitors in the form of SFRPs are required to ensure that elongation occurs stably along a single axis.

## Discussion

Together, our findings show that an excitable system composed of WNT and FGF signaling drives human axial elongation. A key constituent of this excitable system are NMPs that act as signaling centers in the tailbud. Despite being bipotent, in our organoids NMP-like cells give rise only to neural progenitors, suggesting that their ability to act as signaling centers is independent of their potency to generate mesodermal tissue. Accordingly, organoids consist only of a neural tube and tailbud, and elongate axially in the absence of any mesoderm. Furthermore, as with any excitable system, organoids are prone to runaway events, leading to ectopic sites of elongation. These runaway events must be dampened by secreted WNT inhibitors to maintain a single axis during human A-P morphogenesis (Fig. 4J).

We anticipate that our model of the tailbud and elongating neural tube as an excitable system could generalize to other organ primordia in which asymmetric signals are coupled to morphogenesis, especially those in which WNT and FGF signaling have essential roles. These include the limb bud (Zeller et al., 2009) and lung bud (Volckaert and De Langhe, 2015), where polarization and elongation are essential for proper development. Furthermore, the ability of our organoids to robustly generate NMPs in the tailbud opens avenues to studying the coordination of human spinal cord and musculoskeletal development, as we show in a companion manuscript (Yaman et al., 2022). We suggest that by systematically building up the complexity of tissue-tissue interactions present in the embryo, we may uncover developmental mechanisms that are difficult to dissect *in vivo*.

Stochasticity has attracted much attention in biological systems. The large variability exhibited by *in vitro* developmental model systems is often attributed to this intrinsic noise. However, our study shows that much of this variability can be eliminated if the boundary conditions for the underlying dynamical systems can be controlled, in this case by coupling these systems to each other. Learning to reduce variability in biological systems using a combination of bioengineering and machine learning tools will aid in the discovery of mechanisms underlying the development of complex human tissues.

## Acknowledgments

We thank Nicole Ramirez, Claire Reardon, and Zachary Niziolek at the the Bauer Core Facility at Harvard University for single cell RNA sequencing and assistance with FACS, and Doug Richardson at the Harvard Center for Biological Imaging for help with imaging. We thank Perry Ellis, members of the Weitz Lab, and Professor Ian Papautsky for their contributions and advice regarding the microfabrication methods used for this manuscript. We thank Margarete Diaz Cuadros and the Pourquie lab for sharing the H2B-mCherry hiPSC line used for imaging. We thank Professors Richard Losick, Olivier Pourquie, Iftach Nachman and members of the Ramanathan Lab for scientific discussions and comments. This work was supported in part by 1RF1MH123948-01 (SR, JL), 5R01HD100036-02 (SR) and startup funds from Harvard University.

## Author contributions

GMA and SR designed the study. SHM, HCM and GMA performed the data analysis andsimulations. GMA, TW and HCM performed all the experiments with the exception of STARmap. ZL, YH, JL, and XW designed, performed and analyzed the STARmap experiments. GMA, HCM, SHM, and SR wrote the manuscript.

## Data and materials availability

The single cell RNA-seq dataset generated in this study will be made available in the NCBI GEO repository and the NCBI SRA by the time of publication, or in advance of that date upon reasonable request. All other data and code are available upon reasonable request.

## Supplementary Materials

Figs. S1 to S4

Movie S1

Materials and Methods

Supplementary Methods

---

**Fig. S1.**
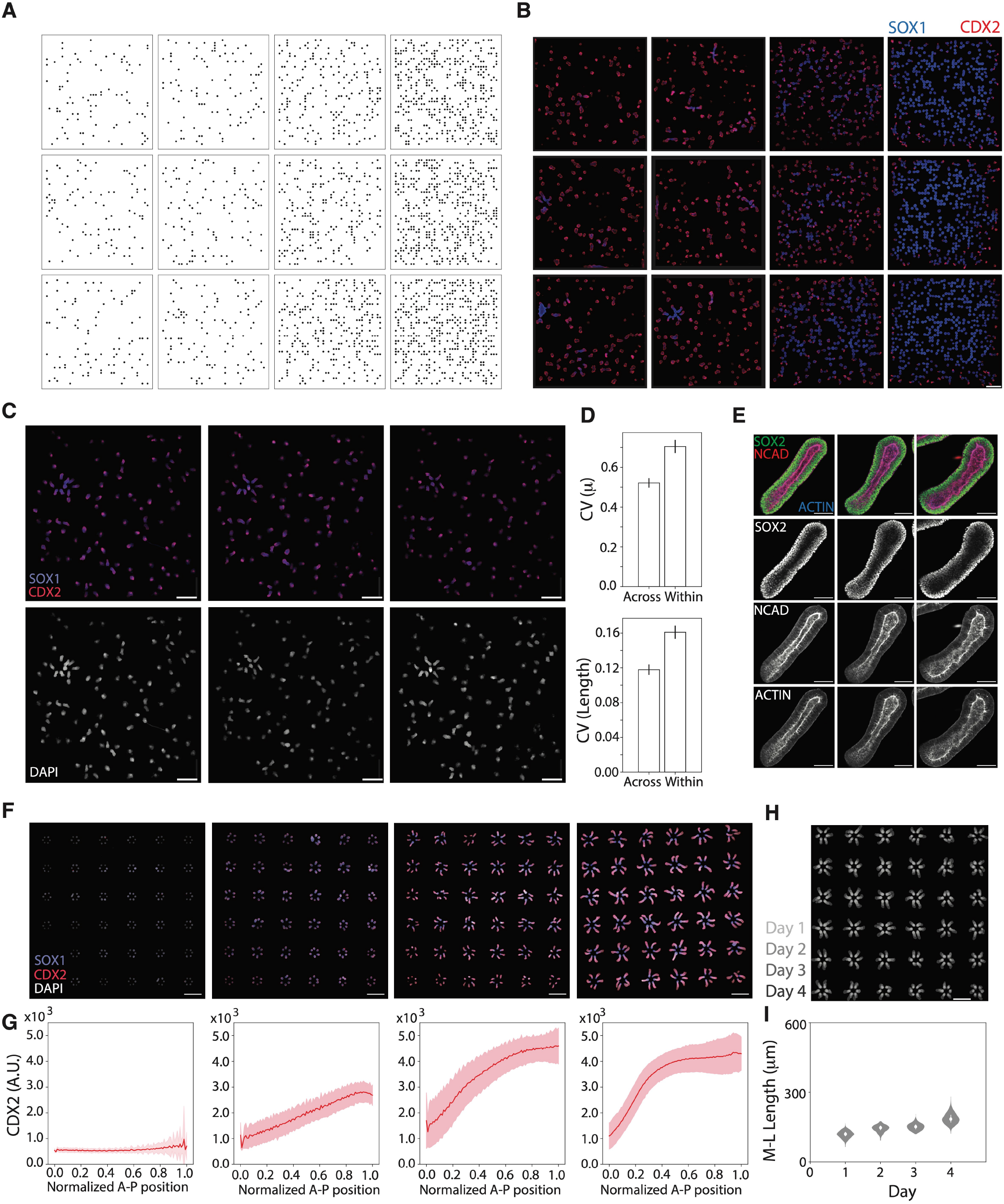
Spatial coupling of organoids influences their axial patterning and morphogenesis. **(A)** Randomly generated micropattern arrays at low density (first and second column), medium density (third column), and high density (fourth column). Micropatterns have a diameter of 150 μm and a minimum spacing of 50 μm (see Supplementary Methods). **(B)** Organoids from random arrays in (A) on day 4, stained for SOX1 (blue) and CDX2 (red). n=2,373 organoids total. Scale bar 1 mm. **(C)** Three replicate experiments (left, middle, right) on a low density random array on day 4, stained for DAPI (top) and SOX1 and CDX2 (bottom). **(D)** Coefficient of variation in measured polarization μ (top) and length based on Feret diameter (bottom) for organoids across and within replicate experiments in (C). Error bars denote SEM (n=105 organoids per experiment). **(E)** Fluorescence confocal images of organoids from hexagon array on day 4 stained for SOX2, N-cadherin (NCAD), and F-actin (ACTIN). Scale bar 100 μm. **(F)** Overlay image of organoid contours in a single experiment from day 1 to day 4. Scale bar 1 mm. **(G)** Organoids on a hexagonal array on consecutive days 1, 2, 3, and 4 stained for DAPI, SOX1, and CDX2. Scale bar 1 mm. **(H)** Plot of CDX2 expression across organoids on hexagon array as a function of normalized A-P position on consecutive days (see Methods). Midline represents mean, shaded region denotes standard deviation (n=216).

**Fig. S2.**
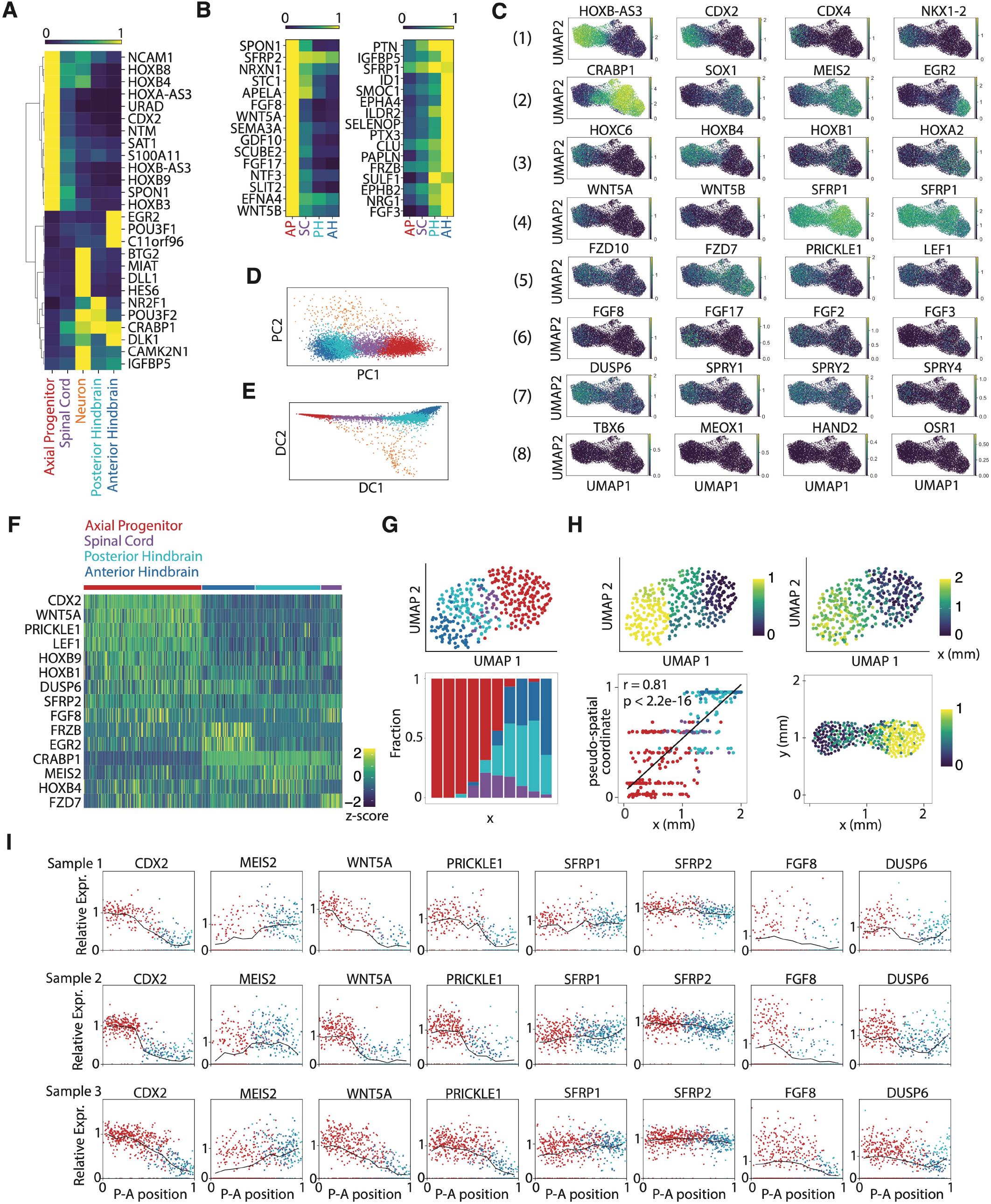
Validation of cell types and signaling profiles in elongating organoids. **(A)** Average gene expression of 26 genes selected by sparse multimodal decomposition (see Methods) across annotated cell types obtained from elongating organoids on day 4 using scRNA-seq. Cell types consist of axial progenitor (CDX2+), spinal cord (HOXB4+ CRABP1+), posterior hindbrain (NR2F1+), anterior hindbrain (EGR2+), and neuronal (HES6+ BTG2+) identities. Expression values are log- and min/max-normalized across clusters. **(B)** Average gene expression of top differentially expressed secreted genes across progenitor clusters, enriched in posterior (left) or anterior (right) cell types (see Methods). **(C)** UMAP plot of cells from scRNA-seq colored by log-normalized gene expression. Rows correspond to different classes of genes, including (1) posterior genes, (2) anterior genes, (3) HOX genes, (4) WNT ligands and inhibitors, (5) WNT pathway genes, (6) FGF ligands, (7) FGF targets, (8) paraxial and lateral mesoderm marker genes. **(D)** PCA plot of cells from scRNA-seq along first two principal components in 26-dimensional gene space, colored by cell type. **(E)**Two-dimensional diffusion map of cells from scRNA-seq, colored by cell type (see Methods). **(F)** Gene expression heatmap of cells (n=359) from organoid sample 1 on day 4 using *in situ* sequencing (STARmap). Cells clustered in a 16-gene space selected based on diffusion map results (see Methods) show axial progenitor (CDX2+ CRABP1-), spinal cord (CDX2+ CRABP1+), posterior hindbrain (CDX2-CRABP1+ EGR2-), and anterior hindbrain (CRABP1+ EGR2+) progenitor identities. Expression values are log- and z-score-normalized across cells. **(G)** *Top:* UMAP plot of cells from STARmap applied to organoid sample 1, colored by cell type. *Bottom:* Fraction of cell types along the elongation axis (x), partitioned into 10 bins of equal length. **(H)** *Top left:* UMAP plot of cells from STARmap applied to organoid sample 1, colored by pseudo-spatial coordinate inferred from DPT (see Methods). **(I)** Spatial P-A gene expression profiles for selected genes from STARmap applied to organoid sample 1 (top, n=359 cells), 2 (middle, n=610 cells), 3 (bottom, n=637 cells) colored by cell type. Each plot displays mean (black) and per-cell (points) log-normalized relative expression values partitioned 10 bins of equal length.

**Fig. S3.**
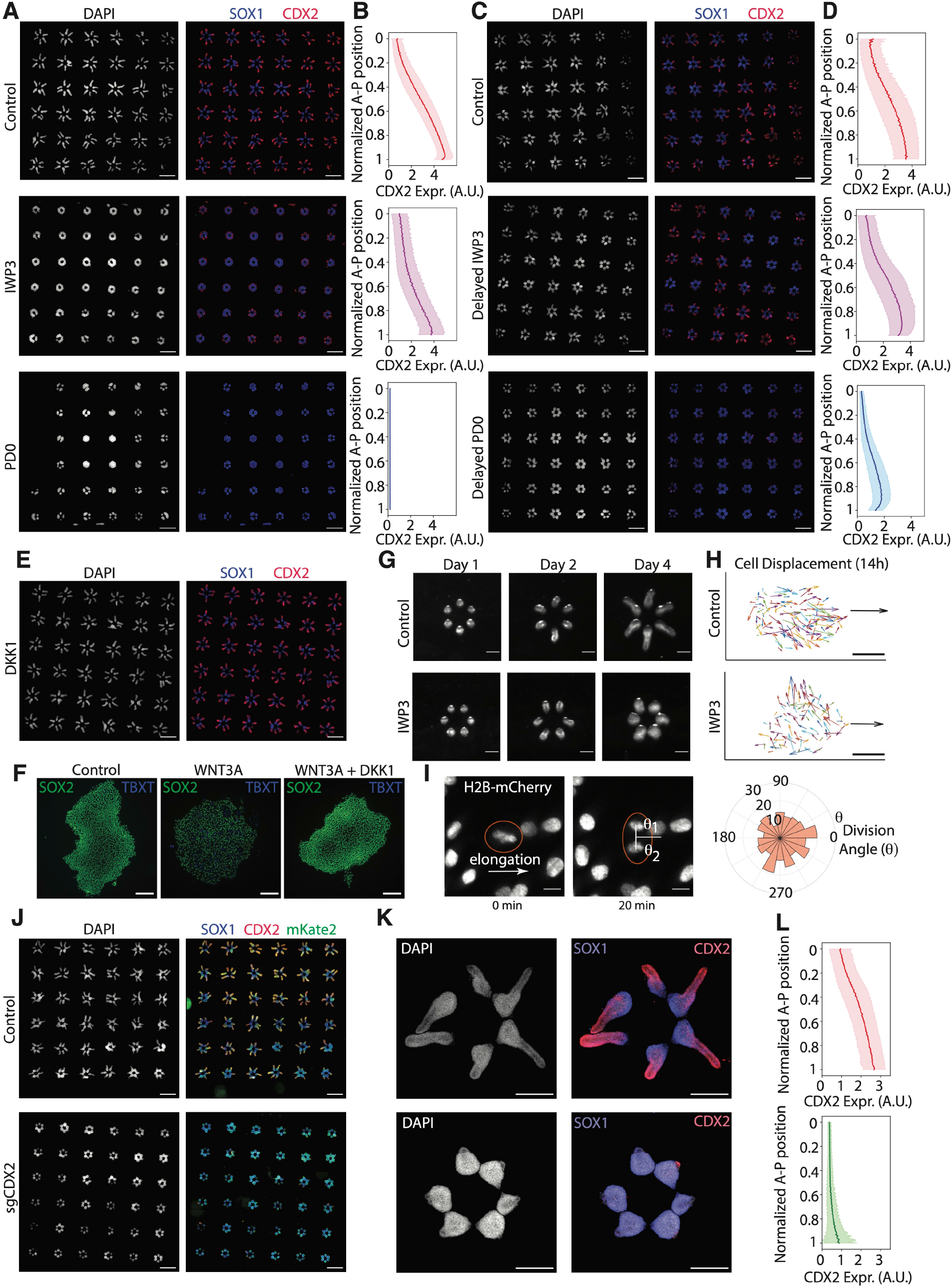
WNT ligands drive elongation downstream of FGF/ERK-dependent CDX2 expression. **(A)** Control (top), IWP3-treated (middle), and PD0-treated (bottom) organoids on day 4 stained for DAPI (left) and SOX1 and CDX2 (right). Scale bar 1 mm. **(B)** Plot of CDX2 expression across control (top, n=211), IWP3-treated (middle, n=176), and PD0-treated (bottom, n=157) organoids as a function of normalized A-P position. Midline represents mean, shaded region denotes standard deviation. **(C)** Control (top), delayed IWP3-treated (middle), and delayed PD0-treated (bottom) organoids on day 4 stained for DAPI (left) and SOX1 and CDX2 (right). Scale bar 1 mm **(D)** Plot of CDX2 expression across control (top, n=212), delayed IWP3-treated (middle, n=214), and delayed PD0-treated (bottom, n=211) organoids as a function of normalized A-P position. Midline represents mean, shaded region denotes standard deviation. **(E)** DKK1-treated organoids on day 4 stained for DAPI (left) and SOX1 and CDX2 (right). **(F)** Control (left), WNT3A-treated (middle), and WNT3A+DKK1-treated (right) hPSC colonies on day 2 stained for SOX2 and TBXT. **(G)** Stereomicroscope images of control (top) and delayed IWP3-treated (bottom) ZO1-EGFP organoids on consecutive days. **(H)** Net cell displacement vectors for tracked cells in control (top) and delayed IWP3-treated (bottom) organoids for 14 hours starting on day 2. Horizontal black arrow indicates direction of axis elongation. Scale bar 100 μm. **(I)** *Left*: Representative live fluorescence images of H2B-mCherry+ cells in control organoids before (left) and after cell (right) division. θ_1_ and θ_2_ mark cell division angles relative to the axis of elongation (horizontal white arrow). Scale bar: 10 μm. *Right*: Histogram of cell division angle relative to the axis of elongation in control organoids over 12 hours starting on day 2 (n=121). **(J)** CRISPRi organoids containing an mKate2-expressing control (top) or CDX2-targeting (bottom, sgCDX2) gRNA constructs on day 4, stained for DAPI (left) and SOX1 and CDX2 (right). Scale bar 1 mm. **(K)** Fluorescence confocal images of CRISPRi organoids carrying control (top) and CDX2-targeting (bottom) gRNA on day 4, stained for DAPI (left) and SOX1 and CDX2 (right). **(L)** Plot of CDX2 expression across control sgRNA (top, n=213) and sgCDX2 (bottom, n=214) CRISPRi organoids as a function of normalized A-P position. Midline represents mean, shaded region denotes standard deviation.

**Fig. S4.**
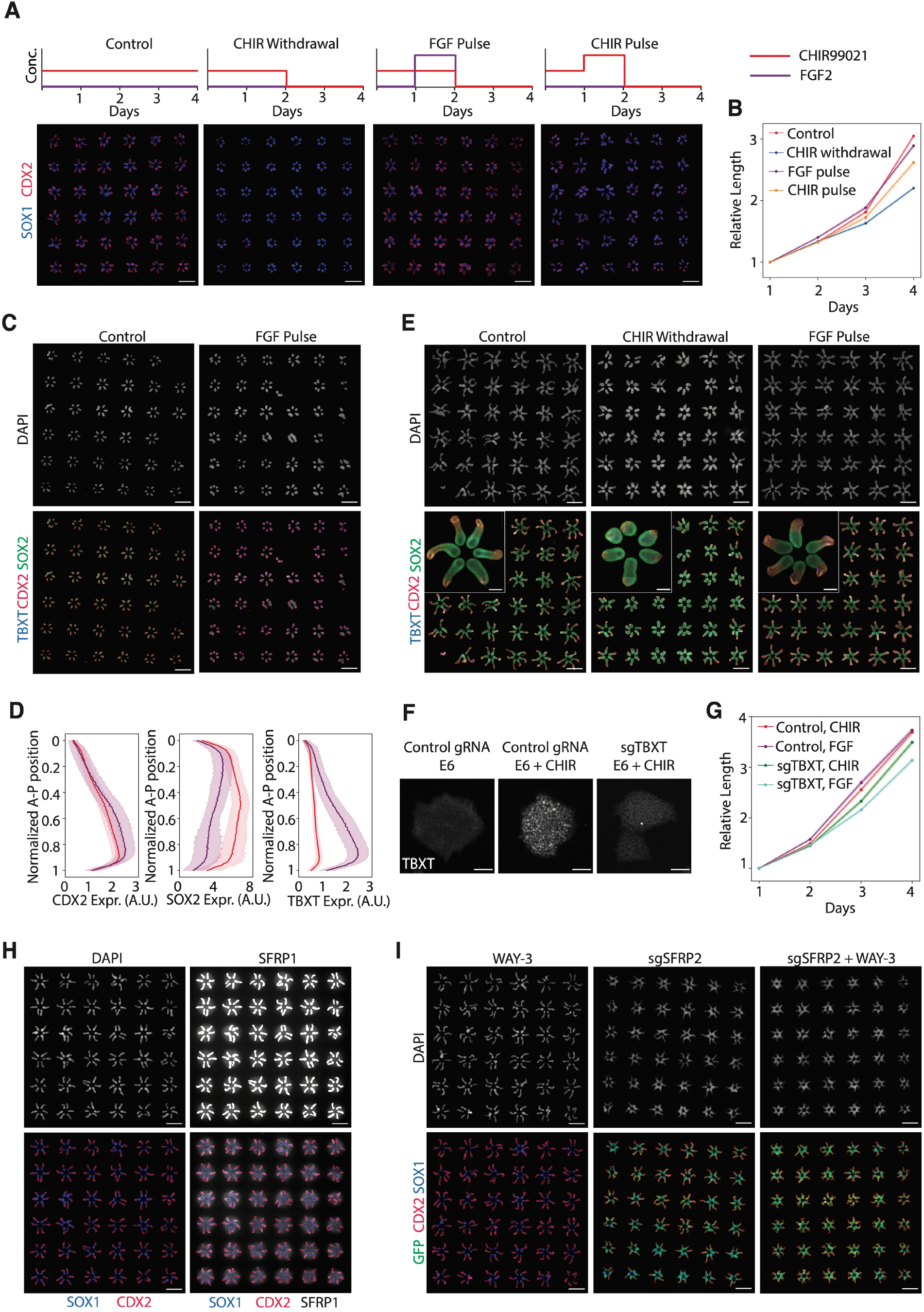
A TBXT-dependent excitable system driving elongation is triggered by induction of an NMP-like signaling center, and is stabilized by secreted WNT inhibitors. **(A)** *Top:* Concentration profiles of CHIR99021 (CHIR) and FGF2 (FGF) in medium for control, CHIR withdrawal, FGF pulse, CHIR pulse conditions. *Bottom:* Control, CHIR withdrawal, FGF pulse, and CHIR pulse-treated organoids on day 4, stained for DAPI (top) and SOX1 and CDX2 (bottom). **(B)** Line plot of relative lengths over time of organoids exposed to control (red), CHIR withdrawal (blue), and FGF pulse (purple) and CHIR pulse (orange) conditions. Shaded regions denote SEM (Control, n = 216, CHIR withdrawal, n = 214, FGF pulse, n = 216, CHIR pulse, n = 214). **(C)** Control (left) and FGF pulse-treated (right) organoids on day 2, stained for DAPI, CDX2, SOX2, and TBXT. Scale bar 1 mm. **(D)** Plots of CDX2 (left), SOX2 (middle), TBXT (right) expression across control (red, n=199) and FGF pulse-treated (purple, n=184) organoids as a function of normalized A-P position. Midline represents mean, shaded region denotes standard deviation. **(E)** Control (left), CHIR withdrawal (middle), and FGF pulse-treated (right) organoids on day 4, stained for DAPI, CDX2, SOX2, and TBXT. Scale bar 1 mm. Inset, *top left*: Zoomed-in views of organoids on day 4 shows retention of TBXT+ cells in FGF pulse condition. Scale bar 200 μm. **(F)** CRISPRi hPSC colonies containing a control gRNA (left, middle) or sgTBXT gRNA construct (right) after 24 hours of exposure to E6 (left) or E6 + CHIR (middle, right) stained for TBXT. Scale bar 200 μm. **(G)** Line plot of relative lengths over time of control sgRNA and sgTBXT CRISPRi organoids exposed to CHIR (control, red; sgTBXT, green) and FGF pulse (control, purple; sgTBXT, turquoise) conditions. Shaded regions denote SEM (Control, CHIR, n = 216; Control, FGF pulse, n = 216; sgTBXT, CHIR, n = 216; sgTBXT, FGF pulse, n = 206). **(H)** Control organoids on day 4, stained for DAPI, CDX2, SOX1, and SFRP1. Images were contrast-adjusted to visualize extracellular SFRP1. In the merged image, DAPI was subtracted to emphasize extracellular SFRP1. Scale bar 1 mm. **(I)** Wild type organoids treated with WAY-3 (left), CRISPRi organoids containing a GFP-expressing guide RNA construct against SFRP2 (sgSFRP2) (middle), and sgSFRP2 CRISPRi organoids treated with WAY-3 (right) on day 4 stained for CDX2, SOX1, and DAPI. Scale bar 1 mm.

**Movie S1.**
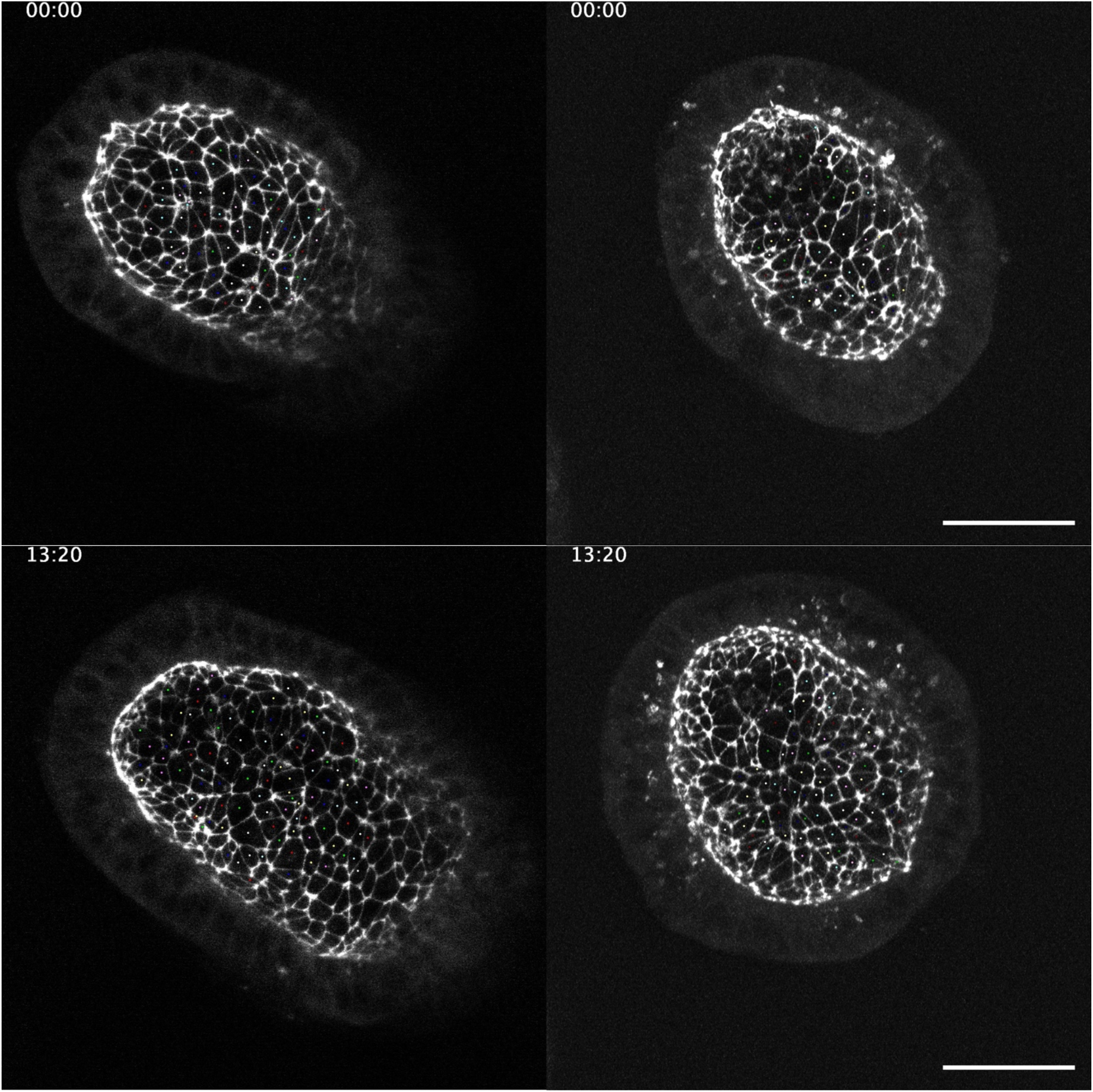
Time-lapse microscopy of ZO1-EGFP+ organoids. Max intensity projections of control (left) and IWP3-treated (right) organoids imaged on an LSM 980 confocal microscope starting on day 2 of differentiation, with z-stacks acquired every 20 minutes using a 488 nm laser (see Methods). The centroid of each cell was tracked manually in FIJI (see Supplementary Methods), and is labeled as a colored dot. Time stamps are in hh:mm format. Scale bar 50 μm.

